# Atypical cell death in the growth plate chondrocytes of *Tric-b*-knockout mice

**DOI:** 10.1101/2023.02.28.530410

**Authors:** Atsuhiko Ichimura, Yuu Miyazaki, Hiroki Nagatomo, Masato Tomizawa, Takaaki Kawabe, Nobuhisa Nakajima, Naoki Okamoto, Shinji Komazaki, Sho Kakizawa, Miyuki Nishi, Hiroshi Takeshima

## Abstract

TRIC-A and TRIC-B proteins form homotrimeric cation-permeable channels in the sarco/endoplasmic reticulum (SR/ER) and nuclear membranes and are thought to contribute to counterionic flux coupled with store Ca^2+^ release in various cell types. Serious mutations in the *TRIC-B* locus cause autosomal recessive osteogenesis imperfecta (OI), which is characterized by insufficient bone mineralization. We have reported that *Tric-b*-knockout mice can be used as an OI model. Here we report irregular cell death in proliferating growth plate chondrocytes in developing *Tric-b*-knockout bones. In the knockout chondrocytes, excess pro-collagen fibers were occasionally accumulated in severely dilated ER elements. Of the major ER stress pathways, the PERK pathway was preferentially hyperactivated in the knockout chondrocytes, and most likely altered gene expression to induce apoptosis-related proteins including CHOP and caspase 12. In Ca^2+^ imaging experiments, the knockout chondrocytes exhibited aberrant Ca^2+^ handling; ER Ca^2+^ release was impaired, and intracellular Ca^2+^ concentration was elevated. Our data suggest that *Tric-b* deficiency directs growth plate chondrocytes to pro-apoptotic stages by compromising cellular Ca^2+^-handling and exacerbating ER stress, leading to atypical apoptotic cell death.

## Introduction

Ca^2+^ store functions organized in the sarco/endoplasmic reticulum (SR/ER) are essential for cellular homeostasis. Store Ca^2+^ fluxes are predominantly mediated by SR/ER Ca^2+^-ATPase pumps and Ca^2+^ release channels, namely inositol trisphosphate (IP_3_R) and ryanodine receptors (RyR), and supposed to accompany counterion fluxes to balance charge, osmolality and pH between the SR/ER lumen and the cytoplasm (Fink & Veigel, 1996, Meissner, 1983, Somlyo et al., 1981). Although several ionic fluxes, such as K^+^, Cl^−^ and H^+^ currents, have been detected in SR/ER membranes (Coronado & Miller, 1980, Ide et al., 1991, Kamp et al., 1998), the molecular basis of the counterionic fluxes is largely unknown. It is reasonably proposed that counterion species and their current densities during Ca^2+^ uptake and release are divergent among cell types, because SR/ER-resident channels and transporters may be differentially expressed to generate various counterion fluxes. We previously identified two <u>tr</u>imeric intracellular <u>c</u>ation channel subtypes, namely TRIC-A and TRIC-B, both of which are distributed to the SR/ER and nuclear membranes and form ionic channels that are predominantly permeable to monovalent cations in planner lipid bilayer membranes (Pitt et al., 2010, Venturi et al., 2013, Yazawa et al., 2007). The unique three-dimensional structures of TRIC channels have been elucidated, and each subunit possesses its own ion-conducting pore equipped with phospholipids under intracellular conditions (Kasuya et al., 2016, Wang et al., 2019, Yang et al., 2016). In knockout mice lacking TRIC subtypes, cell types developing functional defects commonly exhibit impaired Ca^2+^ release and store Ca^2+^ overloading. For example, *Tric-a*-knockout mice exhibit impaired RyR-mediated Ca^2+^ release in muscle cells (Yamazaki et al., 2011, Zhao et al., 2010). In contrast, IP_3_R-mediated Ca^2+^ release is compromised in alveolar epithelial cells and osteoblasts from *Tric-b*-knockout mice (Yamazaki et al., 2009, Zhao et al., 2016). These observations indicate that TRIC channels generate counter-K^+^ fluxes at least in part to facilitate store Ca^2+^ release in various cells. Additionally, more recent observations suggest that TRIC-A directly interacts with and activates RyR channels in addition to providing a counterion current (Zhou et al., 2020).

Long bones develop through the biological process called endochondral ossification, and the initial stage of this process is cartilage formation (Berendsen & Olsen, 2015). During cartilage formation in early embryogenesis, chondroblasts undergo differentiation and organize morphologically distinct zones, each of which contains homogeneous chondrocytes specified by morphological characteristics. Round-shaped chondrocytes propagate in the epiphyseal end and produce type II collagen. Then, the round chondrocytes structurally change into flat chondrocytes that proliferate to arrange characteristic columnar arrays. The columnar chondrocytes subsequently differentiate into hypertrophic chondrocytes expressing type X collagen, and finally swell and undergo apoptosis. Finally, the region scattered with the generated apoptotic bodies is gradually replaced by trabecular bone through the action of osteoclasts and osteoblasts.

Osteogenesis imperfecta (OI) is a genetic disease characterized by repeated bone fractures due to reduced bone mass (Byers & Pyott, 2012). The majority of OI cases result from defective type I collagen; structural mutations and altered posttranslational modifications lead to its insufficient synthesis, unfolding, mistrafficking, poor secretion and disincorporation into the bone matrix. OI-causing mutations are also found in collagen-unrelated genes, such as the osteoblast-specific transcription factor Osterix and the osteoblast-specific transmembrane protein IFITM5. Furthermore, homozygous deletion mutations in the *TRIC-B* (also referred to as *TMEM38B*) locus have been identified in several OI pedigrees (Lv et al., 2016, Rubinato et al., 2014, Shaheen et al., 2012, Volodarsky et al., 2013); the critical mutations are all supposed to produce defective TRIC-B channels in the reported patients. Indeed, *Tric-b*-knockout mice develop an OI-like phenotype, and the *Tric-b* deficiency induces store Ca^2+^ overloading due to compromised Ca^2+^ release in osteoblasts (Zhao et al., 2016). The experimental evidence indicates that the pro-collagen processing in the ER is likely deranged by the defective store Ca^2+^ handling of osteoblasts, leading to insufficient bone matrix and resulting in poor bone mineralization in OI patients with *TRIC-B* mutations (Bateman et al., 2016, Zhao et al., 2016). However, the impact of the *Tric-b* deficiency has not been examined in resident growth plate chondrocytes in developing bones. In this report, we investigated the irregular cell death observed in proliferating growth plate chondrocytes from developing bone in embryonic *Tric-b*-knockout mice.

## Results

### Dead cells in Tric-b-knockout growth plates

We first explored the impact of *Tric-b* deficiency on growth plate chondrocytes by histologically analyzing developing bones from the *Tric-b*-knockout mice just before birth (E18.5). Consistent with the previous observation that the body size of *Tric-b*-knockout mice is marginally decreased when compared with that of wild-type controls (Yamazaki et al., 2009), the femoral length was slightly reduced in the knockout mice (Fig. EV1A). In the longitudinal femoral sections prepared from *Tric-b*-knockout mice, round, columnar and hypertrophic chondrocyte layers were regularly formed to constitute the developing growth plates (Fig. 1A). The knockout and wild-type growth plate chondrocytes were roughly similar in morphology and density, but the cell sizes of round and columnar chondrocytes were slightly increased in the knockout mice (Fig. 1B). Therefore, *Tric-b* deficiency probably affected cellular integrity in proliferating growth plate chondrocytes.

**Figure 1.**
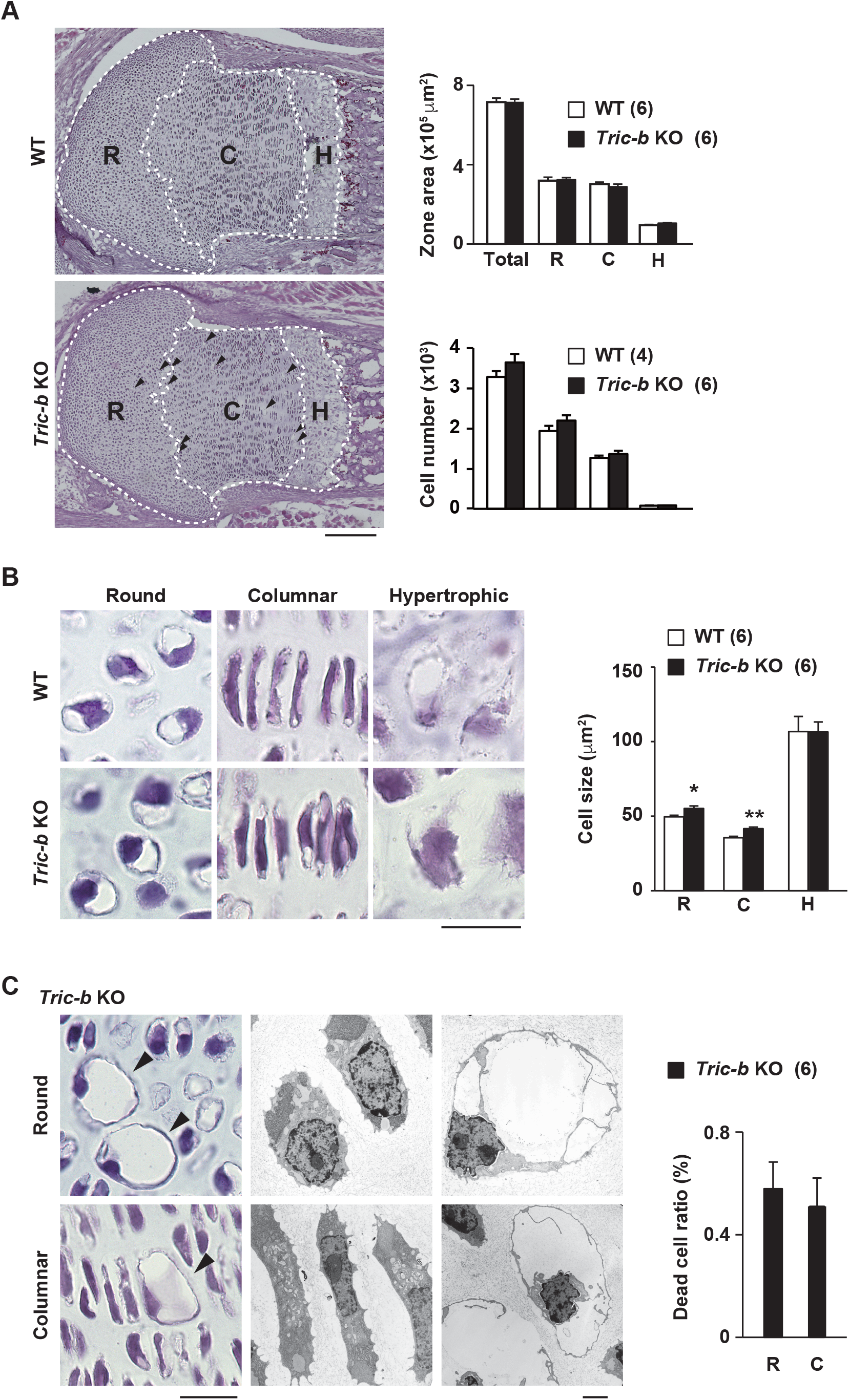
Aberrant cell death in *Tric-b*-knockout growth plates. **A** Regular cell layers formed in *Tric-b*-knockout growth plates. Hematoxylin-eosin-stained histological images of cell layer zones constituted by round (R), columnar (C) and hypertrophic (H) chondrocytes in E17.5 femoral growth plates (left panels; scale bar, 200 *μ*m). Statistical comparison of the zonal sizes and cell numbers in round, columnar and hypertrophic cell layers between *Tric-b*-knockout and wild-type femoral growth plates (bar graphs). **B** Representative photomicroscopic images of hematoxylin-eosin-stained growth plate chondrocytes (left panels; scale bar, 20 *μ*m), and statistical comparison of cell sizes between *Tric-b*-knockout and wild-type growth plate chondrocytes (bar graph). **C** Photo-microscopic images of dilated dead cells (left panels), electron-microscopic images of apparently normal cells (middle panels) and dilated dead cells (right panels) observed in *Tric-b*-knockout round and columnar cell layers. Scale bars, 20 *μ*m. Summarized frequencies of dilated dead round and columnar chondrocytes (bar graph). In the bar graphs, the data are presented as the mean ± SEM., and the numbers of mice examined are shown in parentheses. Statistical differences between the genotypes are indicated by asterisks (\**p*<0.05 and \*\**p*<0.01 in *t*-test).

Unexpectedly, *Tric-b*-knockout growth plates contained severely dilated cells, which were largely negative in hematoxylin-eosin staining and randomly located in the round and columnar cell layers (Fig. 1C). In electron-microscopic observation, the dilated cells contained highly condensed nuclei and almost empty cytoplasm indicative of apoptosis. Although the dilated dead cells were very low in frequency (∼0.6% of total cells in round and columnar cell layers), such irregular cells were never detected in wild-type femoral bones. Furthermore, the dead cells were also detected in the growth plates of humeral and rib bones isolated from the knockout mice (Fig. EV1B). Therefore, *Tric-b* deficiency seemed to occasionally cause apoptotic cell death in proliferating growth plate chondrocytes during long bone development.

### Pro-collagen accumulation in Tric-b-knockout chondrocytes

Proliferating growth plate chondrocytes mainly produce type II collagen as a major cartilage matrix component. We next focused on collagen synthesis in *Tric-b*-knockout chondrocytes. In wild-type round chondrocytes from developing femoral bones, pro-collagen deposits occasionally appeared as intracellular puncta (>5 *μ*m^2^) without colocalization with the ER maker KDEL sequence (Fig. 2A upper panels). However, in *Tric-b*-knockout chondrocytes, pro-collagen deposits were more frequently detected and were colocalized with the ER marker (Fig. 2B). Furthermore, the deposits became larger and denser in *Tric-b*-knockout growth plates, and such severely expanded deposits sporadically covered the bulk of cytoplasm in the presumed dying chondrocytes (Fig. 2A lower panels).

**Figure 2.**
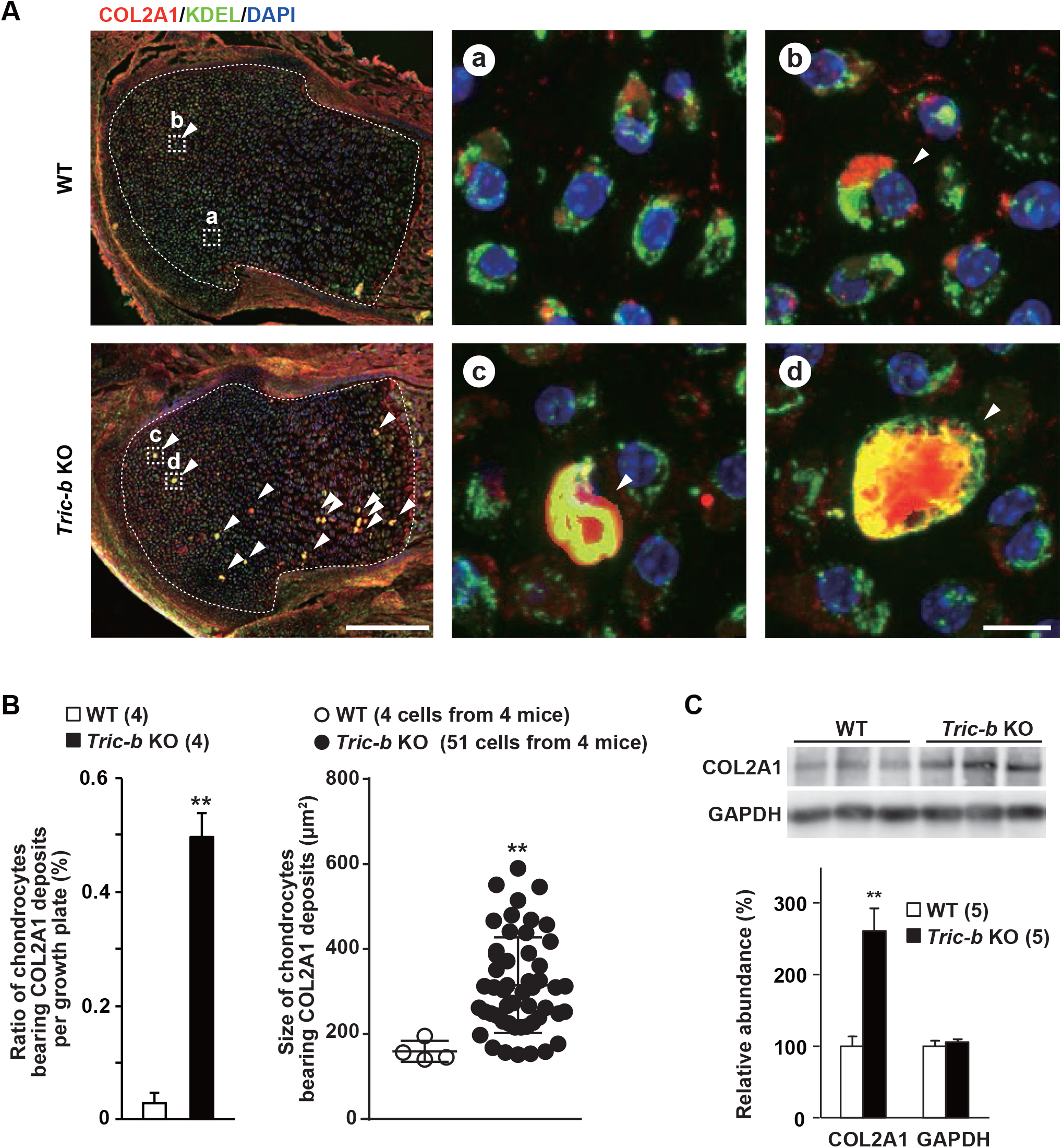
Pro-collagen overaccumulation in *Tric-b*-knockout round chondrocytes. **A** Confocal microscopic images of round chondrocytes, nuclear-stained with DAPI and fluorescence-stained using antibodies against pro-COL2A1 and the ER marker KDEL. As seen in the left low-magnification images (scale bar, 200 *μ*m), pro-COL2A1-positive intracellular deposits (>140 *μ*m^2^, arrowheads) were frequently detected in the round cell layers of *Tric-b*-knockout growth plates. The growth plate chondrocyte zones are indicated by the white dashed lines, and high-magnification images of the cells presenting are indicated by the white dashed boxes. In the high-magnification images (scale bar, 10 *μ*m), wild-type growth plate panels show regular round chondrocytes (a) and one pro-COL2A1 deposit-bearing cell (b), while *Tric-b*-knockout growth plate panels show dilated cells developing pro-COL2A1 deposits (c, d). **B** Appearance ratios and cell sizes of pro-COL2A1-positive chondrocytes in growth plates. **C** Western blot analysis of intracellular pro-COL2A1 in growth plate chondrocytes. Representative COL2A1-immunoreactivities were shown (upper panel), and the digitalized immunoreactivities are statistically compared between wild-type and *Tric-b*-knockout round chondrocytes (lower bar graph). Glyceraldehyde-3-phosphate dehydrogenase (GAPDH) was used as a loading control. In the bar and scatter graphs, the data are presented as the mean ± SEM, and the numbers of mice examined are shown in parentheses. Statistical differences from the genotypes are indicated by asterisks (\*\**p*<0.01 in *t*-test).

To assess the relative amount of pro-collagen within the cells, we solubilized intracellular proteins in a deoxychorate-containing solution and removed extracellular mineralized matrix by centrifugation (Fig. 2C). *Tric-b*-knockout lysates reproducibly exhibited dense immunostaining signals against collagen type II, although signals against the control glycolytic enzyme were similar between wild-type and knockout lysates. The results, together with the histochemical observations, indicated that the ER elements were overloaded with pro-collagen fibers in *Tric-b*-knockout chondrocytes. It is reasonably proposed that *Tric-b* deficiency deranges the ER processing or ER-Golgi trafficking of pro-collagen fibers.

### ER stress in Tric-b-knockout chondrocytes

When immature protein levels reach maximal acceptable levels in the ER lumen, the major transmembrane sensors IRE1 (transmembrane protein kinase inositol-requiring enzyme 1), ATF6 (activating transcription factor 6) and PERK (PKR-like ER kinase) activate the unfolded protein response (UPR) as a homeostatic mechanism in response to ER stresses (Hetz et al., 2020). For example, the ER chaperone BiP/GRP78 is susceptibly induced by activation of either the sensor proteins in various cell types. Under physiological conditions, UPR is moderately activated in cell types that abundantly produce secretory proteins, such as growth plate chondrocytes, pancreatic *β* cells and plasma cells that extensively produce collagen, insulin and antibodies, respectively. To investigate atypical UPR levels in *Tric-b*-knockout chondrocytes, we prepared cell lysates and total RNA from the round cell layers formed in the femoral epiphyses. In Western blot analysis, BiP contents were higher in the knockout lysates than those in wild-type lysates, indicating UPR hyperactivation in *Tric-b*-knockout chondrocytes (Fig. 3A).

**Figure 3.**
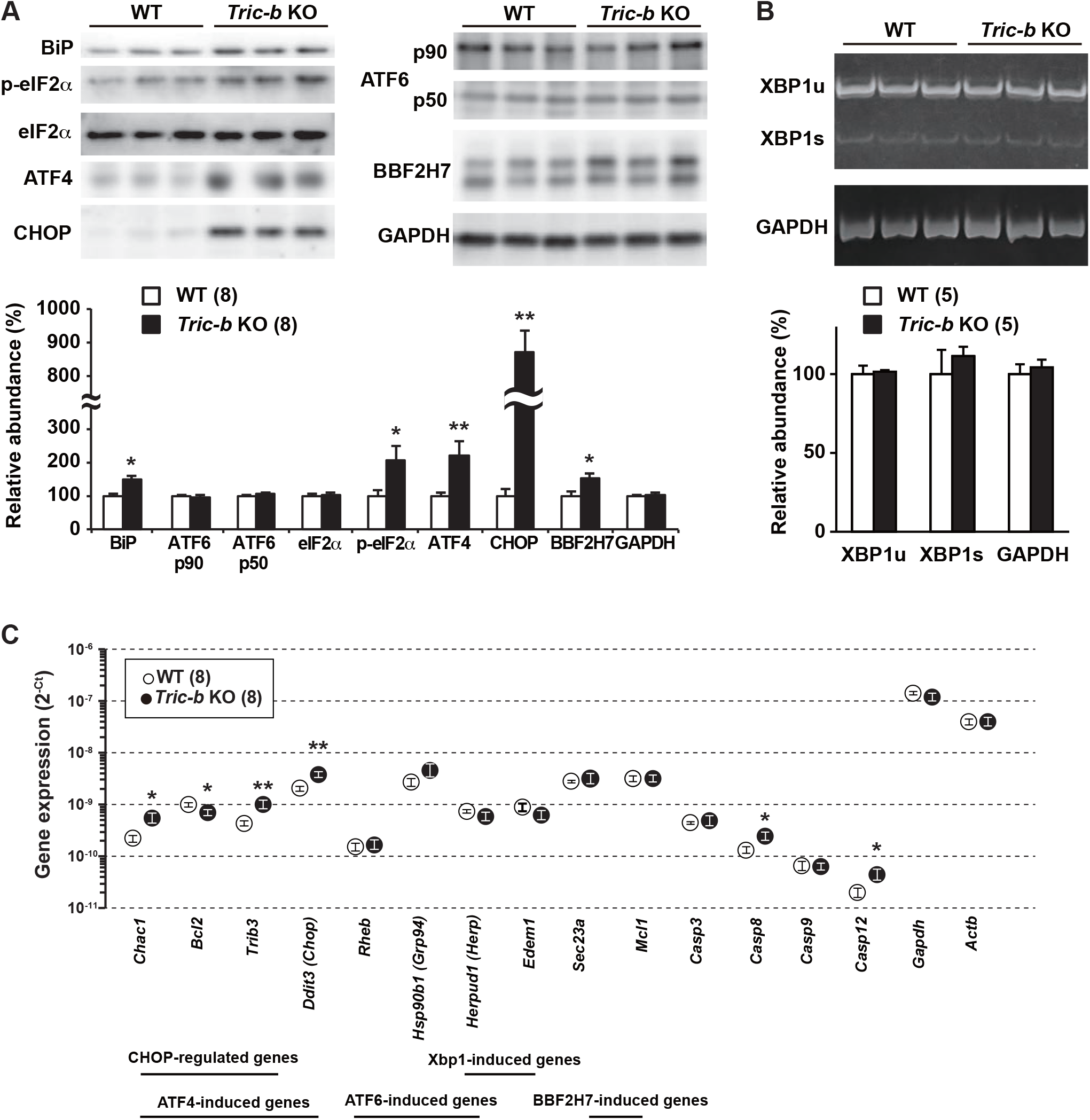
Altered ER stress responses in *Tric-b*-knockout chondrocytes. **A** Western blot analysis of UPR-related proteins in lysates prepared from round chondrocyte-enriched growth plates. Target protein levels were monitored using antibodies against BiP, phospho- and total-eIF2*α*, ATF4, CHOP, p90 and p50 forms of ATF6 and BBF2H7. Representative immunoblot images are shown in the upper panels, and relative immunoreactivities normalized to wild-type values are presented in the bar graph. GAPDH was used as a loading control. **B** RT-PCR analysis monitoring the unspliced (XBP1u) and spliced forms of XBP1 (XBP1s) mRNAs in round chondrocyte-enriched growth plates. Electrophoresis gel images of amplified cDNAs are shown in the upper panel, and relative cDNA intensities normalized to wild-type values are presented in the bar graph. GAPDH mRNA was also amplified as an internal control. **C** Quantitative RT-PCR analysis monitoring mRNAs transcribed from UPR-related genes in round chondrocyte-enriched growth plates. The cycle threshold (*Ct*) indicates the cycle number at which the amount of amplified cDNA reaches a fixed threshold in each RT-PCR reaction. In the graphs, the data are presented as the mean ± SEM, and the numbers of mice examined are shown in parentheses. Statistical differences from the genotypes are indicated by asterisks (\**p*<0.05 and \*\**p*<0.01 in *t*-test).

Activated PERK phosphorylates the translation initiation factor eIF2*α* to reduce translation of newly synthesized proteins and to selectively induce the transcription factor ATF4 (Hetz et al., 2020). ATF4 induces *Chac1* and *Trb3* gene expression (Mungrue et al., 2009). Western blot analysis showed that both phospho-eIF2*α* and ATF4 contents were remarkably elevated in the knockout lysates (Fig. 3A). RT-PCR analysis indicated that *Chac1* and *Trib3* mRNAs were inducibly transcribed in the knockout chondrocytes (Fig. 3C). Therefore, PERK was probably hyperactivated in the knockout chondrocytes.

IRE1 activation stimulates *Xbp1* mRNA splicing and thus promotes the transcription of XBP1-induced genes including *Edem1* and *Erdj4* (Sadighi Akha et al., 2011). In RT-PCR analysis, *Tric-b*-knockout growth plates exhibited no aberrant features in *Xbp1* mRNA splicing and XBP1-induced gene expression (Fig. 3B and 3C). On the other hand, ATF6 p90 is cleaved under ER stress conditions, and the resulting cytoplasmic fragment ATF6 p50 translocates into the nucleus as an active transcription factor to induce the expression of the ATF6-induced genes, including *Grp94* and *Herp* (Kokame et al., 2001). We observed similar ATF6 cleavage and ATF6-induced gene expression between *Tric-b*-knockout and control specimens (Fig. 3B and 3C). Therefore, of the major UPR pathways, IRE1 and ATF6 signaling might be functioning regularly, while PERK pathway seemed to be specifically hyperactivated in *Tric-b*-knockout chondrocytes. This conclusion was further supported by the microarray data derived from round chondrocyte-enriched specimens (Fig. EV2); the heatmap data suggested that CHOP-regulated and ATF4-induced genes were preferentially activated presumably downstream of PERK hyperactivation in the knockout chondrocytes.

BBF2H7 (box B-binding factor 2 human homolog on chromosome 7) is an ER stress sensor essential for chondrogenesis, and its cleaved activation stimulates *Sec23a* and *Mcl1* gene expression (Saito et al., 2009). Sec23a contributes to COPII vesicle formation for ER-Golgi trafficking, while Mcl1 (myeloid cell leukemia sequence 1) belongs to the anti-apoptotic BCL-2 (B-cell leukemia/lymphoma 2) family. Impaired *Sec23a* and *Mcl1* expression promotes ER dilation and apoptosis in *Bbf2h7*-knockout chondrocytes (Saito et al., 2009), similar to the morphological abnormalities observed in *Tric-b*-knockout chondrocytes. Western blotting results showed that BBF2H7 was cleaved more than normal in the knockout chondrocytes (Fig. 3A), and RT-PCR analysis indicated that both *Sec23a* and *Mcl1* genes were similarly activated between the knockout and wild-type chondrocytes (Fig. 3C). Therefore, the BBF2H7 pathway seemed to function normally in *Tric-b*-knockout chondrocytes.

### Apoptosis-related alterations in Tric-b-knockout chondrocytes

It has been reported that caspase 8 (CASP8), CASP9 and CASP12 become differentially active to function as initiator caspases under severe ER stress conditions in several cell types (Kesavardhana et al., 2020). Western blotting suggested no aberrant activation of CASP8 and CASP9 in *Tric-b*-knockout chondrocytes (Fig. 4A). In contrast, both intact and cleaved forms of CASP12 were more abundant in the knockout growth plates than in wild-type controls. Therefore, of the ER stress-related caspase subtypes, CASP12 was most likely preferentially activated in *Tric-b*-knockout chondrocytes.

**Figure 4.**
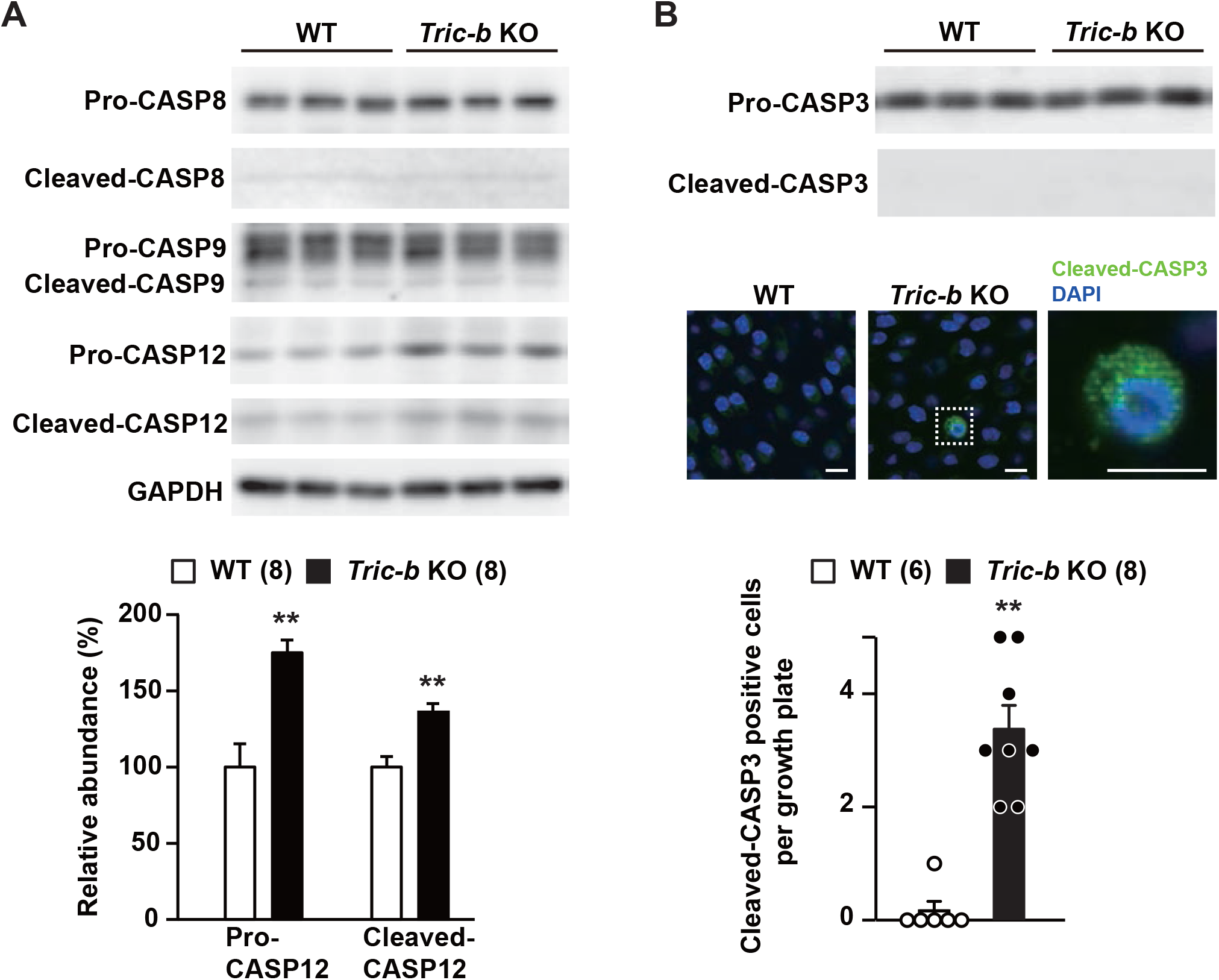
Caspase subtypes in *Tric-b*-knockout chondrocytes. Western blot analysis of CASP8, CASP9 and CASP12 in cell lysates prepared from round chondrocyte-enriched growth plates. Both pro- and cleaved forms of the caspase subtypes were detected by specific antibodies, and representative immunoreactivities are presented (upper panels). The immunoreactivities of pro- and cleaved-CASP12 were digitalized and are statistically compared between wild-type and *Tric-b*-knockout specimens (bar graph). GAPDH was also analyzed as an internal control. **B** Immunochemical analysis of CASP3. Western blot analysis in round chondrocyte-enriched lysates indicated similar pro-CASP3 levels in wild-type and *Tric-b*-knockout specimens, but failed to detect cleaved-CASP3 in both the specimens (upper panels). However, immunohistochemical analysis occasionally detected cleaved-CASP3-positive cells in *Tric-b*-knockout round chondrocyte layers (middle panels, scale bar, 10 *μ*m). The appearance frequencies of cleaved-CASP3-positive cells were statistically compared between wild-type and *Tric-b*-knockout growth plates (bar graph). In the bargraphs, the data are presented as the mean ± SEM., and the numbers of mice examined are shown in parentheses. Statistical differences from the genotypes are indicated by asterisks (\*\**p*<0.01 in *t*-test). ns.: not significant.

CASP3 catalyzes the cleavage of many key cellular proteins and serves as a major effector caspase that executes apoptosis (Kesavardhana et al., 2020). Western blotting failed to detect cleaved-CASP3 in both *Tric-b*-knockout and control growth plates (Fig. 4B). However, immunohistochemical analysis occasionally detected cleaved-CASP3-positive dilated cells in *Tric-b*-knockout growth plates, while such cleaved-CASP3-positive cells were never observed in wild-type growth plates (Fig. 4B). Therefore, the cleaved-CASP3-positive cells were probably assigned as dilated chondrocytes undergoing apoptosis in *Tric-b*-knockout growth plates. Overall, CASP12 and CASP3 likely contributed to *Tric-b*-knockout apoptosis as initiator and effector caspases, respectively.

### Altered store Ca^2+^ handling in Tric-b-knockout chondrocytes

In our previous studies, impaired store Ca^2+^ release and store Ca^2+^ overloading were commonly observed in functionally defective cells prepared from *Tric-a*- and *Tric-b*-knockout mice (Yamazaki et al., 2009, Yamazaki et al., 2011, Yazawa et al., 2007, Zhao et al., 2010). To examine store Ca^2+^ handling of *Tric-b*-knockout chondrocytes, we prepared slice specimens from embryonic femoral bones and collected Fura-2 imaging data from round chondrocytes. Using a perfusion protocol with normal, Ca^2+^-free, ATP-supplemented and Ca^2+^ ionophore ionomycin-containing bathing solutions (Fig. 5A), we examined IP_3_-induced Ca^2+^ release in response to purinergic P2Y receptor activation, ionomycin-induced Ca^2+^ leak and store-operated Ca^2+^ entry (SOCE). In *Tric-b*-knockout chondrocytes, Ca^2+^ transients evoked by P2Y receptor activation became weak, but ionomycin-induced Ca^2+^ leak and SOCE remained unaltered (Fig. 5A). Therefore, IP_3_-induced Ca^2+^ release was significantly impaired in the knockout chondrocytes. However, it was rather surprising that intracellular stores were not Ca^2+^-overloaded despite the impaired IP_3_R-mediated Ca^2+^ release in the knockout chondrocytes (Fig. 5B).

**Figure 5.**
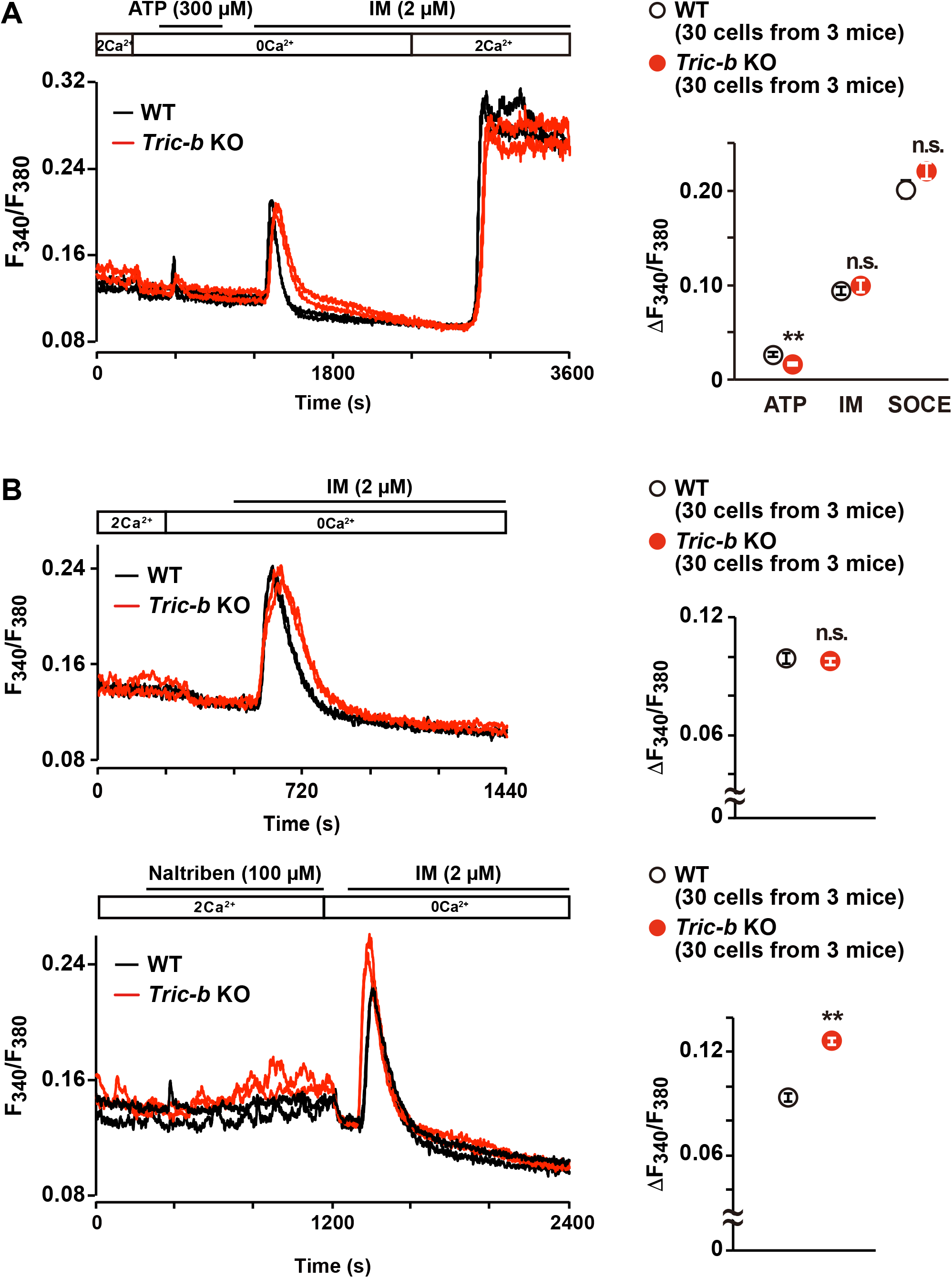
Impaired store Ca^2+^ handling in *Tric-b*-knockout chondrocytes. Round chondrocytes in E17.5 femoral bone slices were examined by Fura-2 imaging. **A** Using the perfusion protocol indicated, representative recording traces obtained from two cells in wild-type (WT) and *Tric-b* knockout mice are shown (left panel). ATP-evoked Ca^2+^ transients (ATP), ionomycin-induced Ca^2+^ leak responses (IM) and store-operated Ca^2+^ entry responses (SOCE) are statistically analyzed (dot graph). Note that the elevated resting [Ca^2+^]_i_ observed in the knockout cells is examined n Figs 6 and 7. **B** Ionomycin-induced Ca^2+^ responses with or without naltriben pretreatment. Representative recording traces from two cells in each genotype are shown (left panels), and the observed Ca^2+^ responses were statistically analyzed (dot graphs). The data are presented as the mean ± SEM, and the numbers of cells and mice examined are shown in parentheses. Statistical differences between the genotypes are marked with asterisks (\*\**p*<0.01 in *t*-test). ns.: not significant.

Growth plate chondrocytes generate spontaneous Ca^2+^ influx by intermissive gating of cell-surface TRPM7 channels, which can be pharmacologically activated with the channel agonist naltriben (Qian et al., 2019). Thus, it is presumed that naltriben preconditioning facilitates autonomic Ca^2+^ entry and ensures full Ca^2+^ loading in intracellular stores. Immediately after naltriben pretreatment, ionomycin-induced Ca^2+^ release was obviously higher in *Tric-b*-knockout chondrocytes than in wild-type controls (Fig. 5C). Therefore, *Tric-b* deficiency might decelerate Ca^2+^ leakage from full stores, thus generating temporal Ca^2+^-overloaded stores. The temporal Ca^2+^ overloading and weakened ATP-induced Ca^2+^ release were consistent with the prevailing notion that TRIC channels facilitate Ca^2+^ release by providing counter-cationic currents (Yazawa et al., 2007). It might be a reasonable speculation that IP_3_R gating is enhanced, thus facilitating Ca^2+^ leakage and preventing store Ca^2+^ overloading in the knockout chondrocytes, because steady-state phospholipase C (PLC) activity was likely elevated under PERK-hyperactivated conditions (see below section).

### Facilitated Ca^2+^ entry in Tric-b-knockout chondrocytes

The elevation of resting intracellular Ca^2+^ concentration ([Ca^2+^]_i_) is generally associated with enhanced Ca^2+^ influx. In growth plate chondrocytes, TRPM7 channels are intermittently activated by intrinsic phosphoinositol turnover and predominantly responsible for resting Ca^2+^ influx (Qian et al., 2019). The resting [Ca^2+^]_i_ of *Tric-b*-knockout chondrocytes was significantly elevated in a normal bathing solution (Figs 5A and 6A). However, under Ca^2+^-free, TRPM7 inhibitor FTY720-treated and PLC inhibitor U73122-supplemented conditions, *Tric-b*-knockout and wild-type chondrocytes exhibited similar resting [Ca^2+^]_i_ levels (Fig. 6A-C). Therefore, steady-state PLC activity was probably enhanced, and thus, TRPM7-mediated Ca^2+^ entry was facilitated in *Tric-b*-knockout chondrocytes.

**Figure 6.**
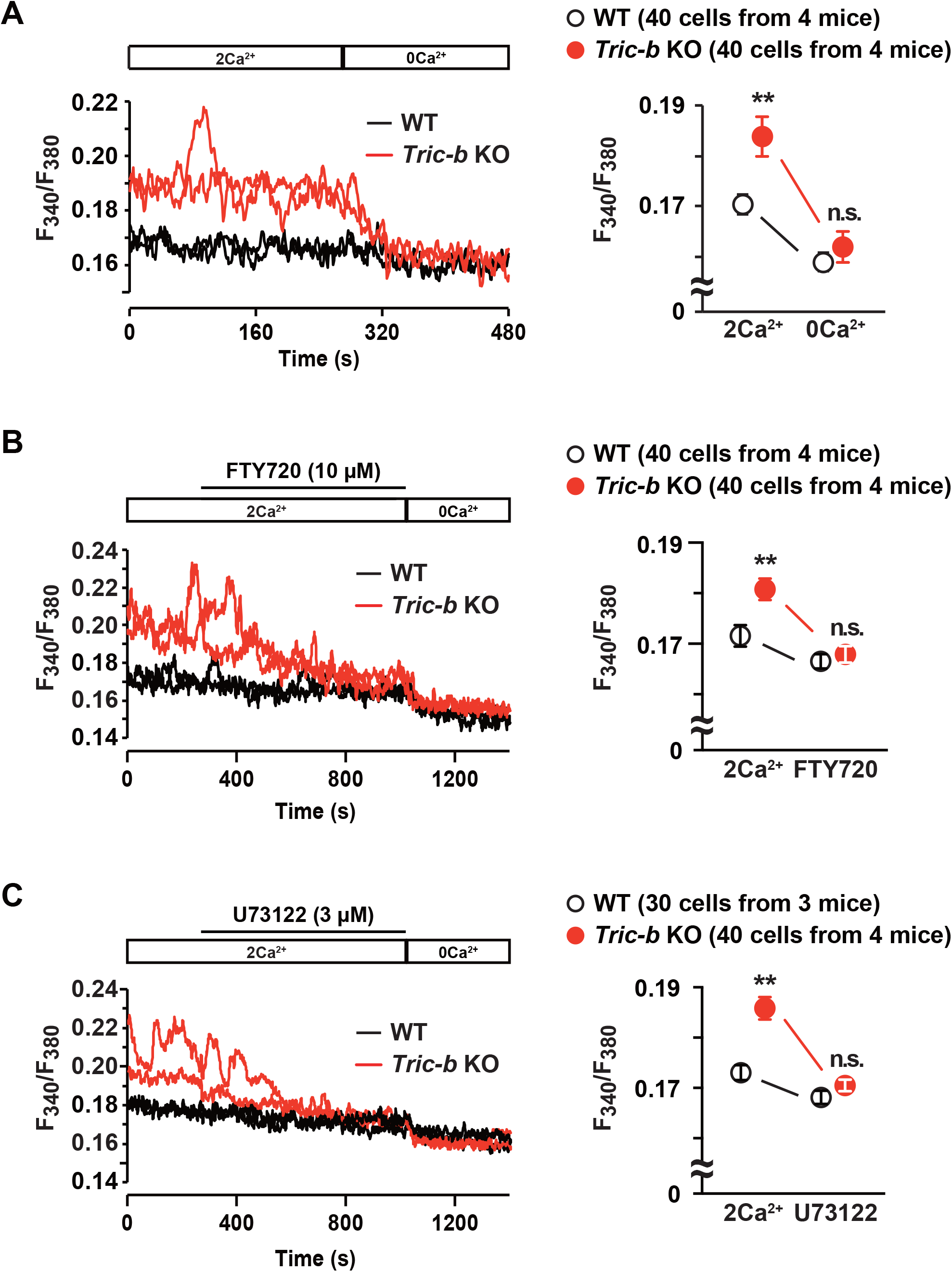
Elevated resting [Ca^2+^]_i_ in *Tric-b*-knockout chondrocytes. In the Fura-2 imaging experiments using the perfusion protocol indicated, representative recording traces obtained from two cells in wild-type (WT) and *Tric-b* knockout mice are shown (left panels). The effects of Ca^2+^-free bathing solution (**A**), the TRPM7 inhibitor FTY720 (**B**) and the PLC inhibitor U73122 (**C**) on resting [Ca^2+^]_i_ are summarized (dot graphs). The data are presented as the mean ± SEM, and the numbers of cells and mice examined are shown in parentheses. Statistical differences between the genotypes are marked with asterisks (\*\**p*<0.01 in *t*-test). n.s.: not significant.

To evaluate the link between hyperactivated PERK and elevated resting [Ca^2+^]_i_ in *Tric-b*-knockout chondrocytes, we utilized the PERK inhibitor GSK2606414 and the PERK activator CCT020312. GSK2606414 treatments (20 *μ*M) did not significantly affect resting [Ca^2+^]_i_ in wild-type chondrocytes but clearly decreased [Ca^2+^]_i_ in the knockout chondrocytes (Fig. 7A). Therefore, under GSK2606414-treated conditions, the knockout and wild-type cells exhibited similar resting [Ca^2+^]_i_. In contrast, CCT020312 treatments (1 *μ*M) obviously elevated [Ca^2+^]_i_ in wild-type chondrocytes but not in the knockout chondrocytes (Fig. 7B). The observations likely suggested that hyperactivated PERK mainly contributed to TRPM7 channel facilitation by stimulating steady-state phosphoinositol turnover in the knockout chondrocytes.

**Figure 7.**
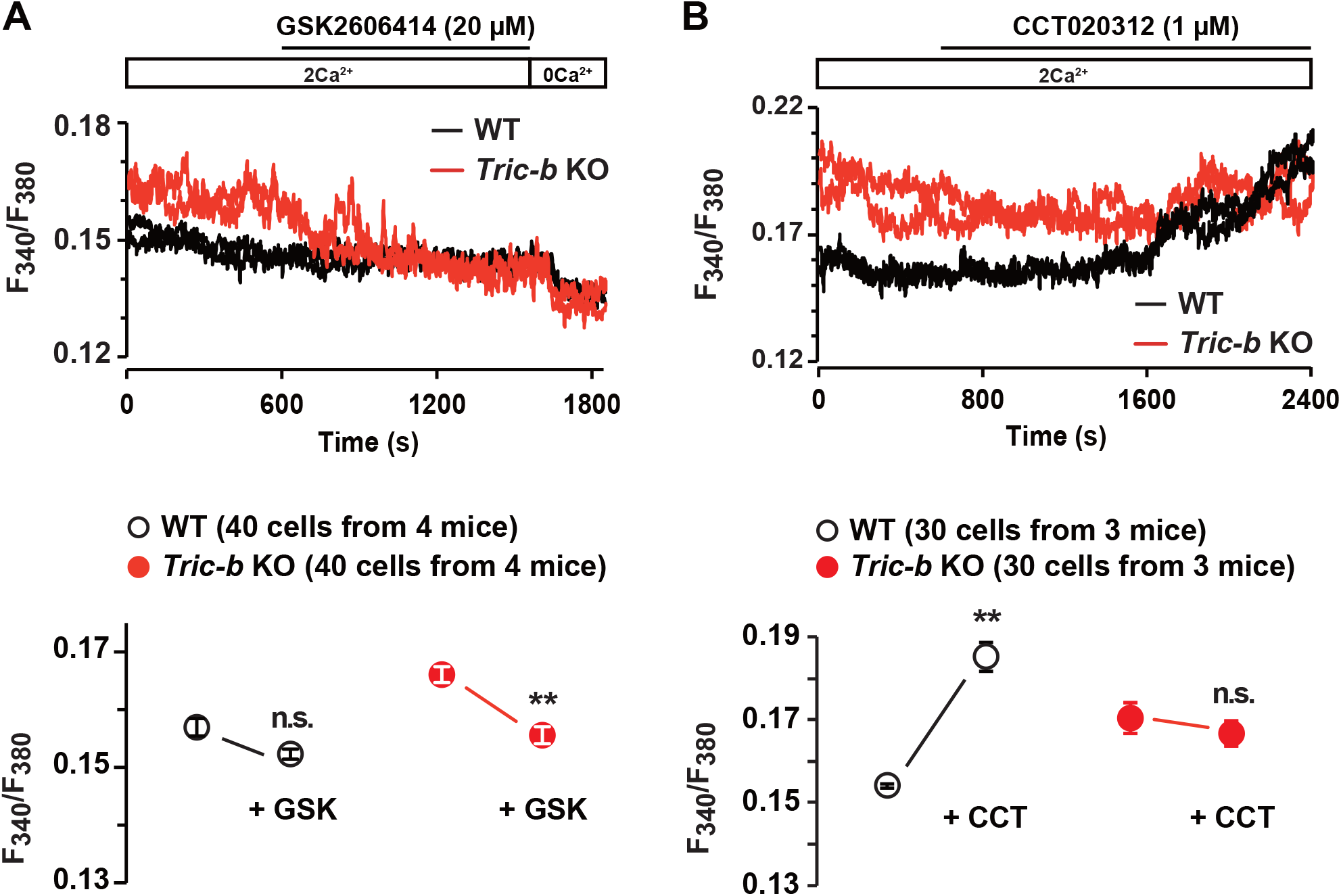
Link between resting [Ca^2+^]_i_ and PERK in *Tric-b*-knockout chondrocytes. In the Fura-2 imaging experiments using the perfusion protocol indicated, representative recording traces obtained from two cells in wild-type (WT) and *Tric-b* knockout mice are shown (upper panels). The effects of the PERK inhibitor GSK2606414 (**A**) and the PERK activator CCT020312 (**B**) on resting [Ca^2+^]_i_ are summarized (dot graphs). The data are presented as the mean ± SEM, and the numbers of cells and mice examined are shown in parentheses. Statistical differences between resting fura-2 ratios before and after drug treatments are marked with asterisks (\*\**p*<0.01 in *t*-test). n.s.: not significant.

## Discussion

Ca^2+^-dependent chaperones and processing enzymes contribute to the maturation of secretory proteins in the ER and Golgi (Lupoli et al., 2018), and vesicular trafficking between the ER and Golgi requires Ca^2+^-dependent processes (Sargeant & Hay, 2022). Defective store Ca^2+^ handling generally aggravates ER stress by disturbing protein processing and vesicular trafficking, and occasionally triggers apoptotic cell death in various cell types. Based on our observations in this study, we propose that *Tric-b* deficiency compromises store Ca^2+^ handling (Fig. 5) and likely deranges processing machinery for secretory proteins, leading to pro-collagen overaccumulation within ER elements (Fig. 2) and PERK hyperactivation (Fig. 3) in growth plate chondrocytes. Furthermore, hyperactivated PERK seems to induce CHOP and Casp12 expression (Fig. 4) and to contribute to PLC activation for facilitating TRPM7-mediated Ca^2+^ entry (Fig. 7) in *Tric-b*-knockout chondrocytes. Therefore, the knockout chondrocytes probably tend to be highly sensitive to apoptosis, because the calpain-Casp12-Casp3 cascade has been reported in various types of apoptosis (Hartl et al., 2009, Lee et al., 2019, Sadasivan et al., 2006). In the knockout chondrocytes that exhibited Casp12 over-expression and high resting [Ca^2+^]_i_, further incidental Ca^2+^ elevation may be predisposed to activate Ca^2+^-dependent calpain for Casp12-cleaved activation, thus triggering the apoptotic cascade. OI patients with *TRIC-B* mutations commonly display short stature (Lv et al., 2016, Rubinato et al., 2014, Shaheen et al., 2012, Volodarsky et al., 2013), and *Tric-b*-deficient mice are small in body size (Yamazaki et al., 2009). Long-bone outgrowth primarily depends on vital proliferation and matrix protein synthesis in growth plate chondrocytes (Berendsen & Olsen, 2015). Perhaps, the growth disturbance observed in OI patients and knockout mice may be primarily caused by the impaired collagen synthesis in proliferating growth plate chondrocytes.

In conventional processing, the three major ER stress sensors PERK, ATF6 and IRE1 become active by the dissociation of the ER chaperone BiP/GRP78 on the luminal sides because excess unfolded proteins attract the chaperone (Hetz et al., 2020). In our observations, PERK was hyperactivated in *Tric-b*-knockout chondrocytes, while ATF6 and IRE1 activation seemed to be similar between the knockout and wild-type cells (Fig. 3). Thus, it is rather mysterious why PERK was specifically hyperactivated in the knockout chondrocytes. Of the three major ER stress pathways, the PERK signaling has been repeatedly reported to be linked with aberrant cellular Ca^2+^ handling (DuRose et al., 2006, Liang et al., 2005, Xu et al., 2017, Zhang et al., 2019, Zhu et al., 2016). For example, the PERK pathway is sensitively activated in response to ER Ca^2+^ depletion (DuRose et al., 2006, Liang et al., 2005), and elevated [Ca^2+^]_i_ activates PERK pathway to promote apoptosis in virus-infected cells (Zhang et al., 2019). Such observations may imply that deranged cellular Ca^2+^-handling preferentially induces PERK activation. Furthermore, in our Ca^2+^ imaging experiments, the PERK inhibitor seemed to immediately attenuate activated Ca^2+^ influx in the knockout chondrocytes, suggesting that downstream of hyperactivated PERK signaling, PLC activity is stimulated, leading to enhanced Ca^2+^ influx by facilitating TRPM7 channel gating. Therefore, we can propose a bidirectional link between PERK signaling and cellular Ca^2+^-handling in growth plate chondrocytes; PERK is hyperactivated by deranged store Ca^2+^ handling due to *Tric-b* deficiency, and hyperactivated PERK stimulates Ca^2+^ influx in a PLC-dependent manner. However, in the PERK signaling cascade proposed thus far, Ca^2+^-dependent machinery for PERK activation and Ca^2+^-handling proteins serving as PERK substrates have not been reported. From a biological point of view, it seems to be important to clarify the molecular mechanism underlying the proposed bidirectional link.

## Materials and Methods

### Histological analyses

All experiments in this study were conducted with the approval of the Animal Research Committee according to the regulations on animal experimentation at Kyoto University. *Tric-b* knockout mice were generated and genotyped as described previously (Yazawa et al., 2007).

For histological analysis, the E18.5 femur, humerus and rib bones were fixed in 4% paraformaldehyde, embedded in Super Cryoembedding Medium (Section-lab, Japan), and frozen in liquid nitrogen. Serial cryosections (∼10 *μ*m thickness) were prepared from the fixed specimens and treated with a commercial hematoxylin and eosin solution (Wako Pure Chemical, Japan) for fluorescence microscopic observation (BZ-X710, Keyence Co., Japan).

For immunohistochemical analysis, the femoral cryosections were treated with 1% bovine serum albumin to block nonspecific binding. The cryosections were incubated with primary antibodies against COL2A1 and KDEL, and then were incubated with an AlexaFluor 488-conjugated antibody against goat anti-mouse IgG and an AlexaFluor 555-conjugated antibody against rabbit IgG (Table S1). After DAPI nuclear staining, fluorescence-labeled sections were examined under a microscope (BZ-X710, Keyence Co.). Captured images from hematoxylin/eosin-stained and fluorescence-labeled sections were quantitatively analyzed using BZ-X analyzer (Keyence Co.) and ImageJ (U.S. National Institutes of Health) software.

### Ultrastructural analysis

For electron-microscopic analysis, the femoral bones were fixed in prefixative solution (3% paraformaldehyde, 2.5% glutaraldehyde, 0.1 M sodium cacodylate, pH 7.5) and placed in postfixative solution (0.1% OsO_4_, 0.1 M potassium ferricyanide, 0.1 M sodium cacodylate, pH 7.4) at room temperature. The specimens were dehydrated using ethanol and acetone, and embedded in Epon to prepare thin sections (100∼150 nm thickness) for analysis under a transmission electron microscope (JEM-200CX, JEOL, Japan).

### Gene expression analysis

Total RNA was prepared from mouse tissues using a commercial kit (Isogen, Nippon Gene). RNA preparations from femoral cartilage plate sections enriched with round chondrocytes were reverse-transcribed and analyzed using the GeneChip Mouse Genome 430 2.0 (Affymetrix) according to the manufacturer’s instructions by an outsourcing company (Takara Bio Co., Japan). The obtained data have been uploaded to the National Center for Biotechnology Information–Gene Expression Omnibus (NCBI-GEO) database (www.ncbi.nlm.nih.gov/geo/) under the accession numbers GSE105256 and GSE223776. The array probe intensities were analyzed with the robust multiarray analysis expression algorithm, which represents the log transformation of intensities (background corrected and normalized) from the gene chips (Gautier et al., 2004), and were visualized in the heatmaps.

To further analyze gene expression, mRNA contents were determined by quantitative RT-PCR as described previously (Miyazaki et al., 2022). Total RNA was reverse-transcribed using the ReverTra ACE qPCR-RT Kit (Toyobo), and the resulting cDNA was examined by real-time PCR (LightCycler 480 II, Roche). The cycle threshold (Ct) was determined from the amplification curve as an index for relative mRNA level in each reaction. The RT-PCR primer sets used in this study are listed in Table S2.

### Immunoblot analysis

Round chondrocyte-enriched growth plate parts were isolated from the E18.5 femoral bones and homogenized in lysis buffer containing 4% sodium deoxycholate, 20 mM Tris-HCl (pH 8.8), 100 mM NaF, 10 mM Na_3_PO_4_, 1 mM Na_3_VO_4_, and 20 mM *β*-glycerophosphate. After sonication (Astrason Ultrasonic Processor XL, Misonix), the homogenates were centrifuged (17,000 x *g*, 30 min) to remove tissue debris. After measurement of total protein concentration (BCA Protein Assay Kit, Pierce), the resulting growth plate lysates were subjected to 8-16.5% SDS–polyacrylamide gel electrophoresis, and separated proteins were then transferred to PVDF membranes (polyvinylidene difluoride, Merck Millipore). After treatments with a blocking reagent (Blocking One solution, Nacalai Tesque, Japan), the membranes were incubated with primary antibodies and then incubated with secondary antibodies; antibodies were diluted in Can Get Signal (Toyobo) before the immunoreaction. Immunoreactivity was visualized using a chemiluminescence reagent (GE Healthcare Life Sciences) and an image analyzer (Amersham Imager 600, GE Healthcare Life Sciences) and were quantitatively analyzed using ImageJ software. The antibodies used in this study are listed in Table S1.

### Fura-2 Ca^2+^ imaging

Bone slices were prepared and Ca^2+^ imaging was performed as described previously (Qian et al., 2019). The bathing solution used was HEPES-buffered saline (150 mM NaCl, 4 mM KCl, 1 mM MgCl_2_, 2 mM CaCl_2_, 5.6 mM glucose, and 5 mM HEPES, pH 7.4). For indicator loading, bone slices prepared using a vibratome slicer were placed on glass-bottom dishes (Matsunami, Japan) and incubated in HEPES-buffered saline containing 15 *μ*M Fura-2 AM (Dojindo) for 60 min at 37°C. For ratiometric imaging, excitation wavelengths of 340 and 380 nm were alternately delivered, and an emission wavelength of >510 nm was detected by a cooled electron multiplying charge-coupled device camera (model C9100-13; Hamamatsu Photonics, Japan). The bone slices were mounted on an upright fluorescence microscope (DM6 FS, Leica) carrying a water immersion objective (HCX APO L 40×, Leica).

### Quantification and statistical analysis

All data obtained are presented as the means ± SEM. with n values indicating the number of examined mice or cells. Student *t*-test and ANOVA were used for two-group and multiple group comparisons, respectively (Prism 7, GraphPad Software Inc.): *p*<0.05 was considered to be statistically significant.

## Data Availability

The datasets generated during and/or analysed during the current study are available from the corresponding author on reasonable request.

## Competing Interest Statement

The authors declare no conflict of financial interest.

## Acknowledgments

We thank Mr. Jun Matsushita (Graduate School of Pharmaceutical Sciences, Kyoto University) for mouse *in vitro* fertilization. This work was supported in part by the MEXT/JSPS (KAKENHI Grant Number 21H02663, 20H03802 and 21K19565), Research Support Project for Life Science and Drug Discovery (Basis for Supporting Innovative Drug Discovery and Life Science Research (BINDS)) from AMED (JP22ama121034), Takeda Science Foundation, Kobayashi International Scholarship Foundation, the NAKATOMI Foundation, Vehicle Racing Commemorative Foundation, Mother and Child Health Foundation and Japan Foundation for Applied Enzymology.

## Author Contributions

A.I. and Y.M. are equally contributing first authors. Y.M., H.N., N.O. and A.I. conducted Ca^2+^ imaging analysis. A.I., Y.M., H.N., M.T., T.K. and N.N. conducted biochemical and cell physiological analysis. S. Komazaki conducted electron microscopy analysis. S. Kakizawa and M.N. contributed to the interpretation of the results. A.I., Y.M. and H.T. drafted the manuscript. H.T. oversaw this project.

## Figure legends

**Figure EV1.**
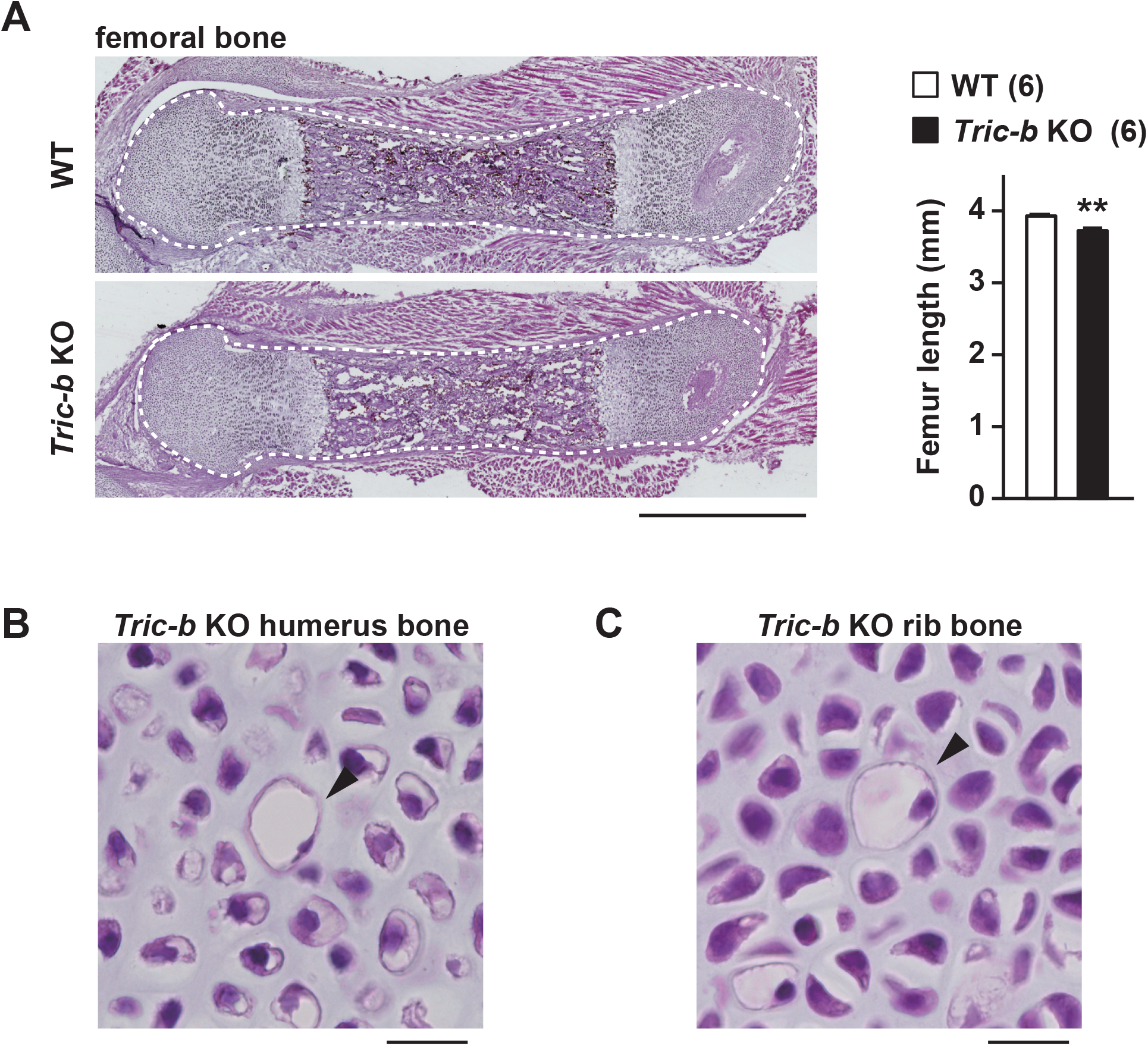
Abnormal histology in the bones of *Tric-b*-knockout mice. **A** Impaired outgrowth in the femoral bones of *Tric-b*-knockout mice. Typical longitudinal femoral sections (hematoxylin-eosin staining) are shown in left panels, and femoral lengths were statistically examined between the knockout and wild-type mice in the bar graph. Scale bar, 1 mm. The data are presented as the mean ± SEM, and the numbers of mice examined are shown in parentheses. Statistical differences between the genotypes are marked with asterisks (\*\**p*<0.01 in *t*-test). **B** Dilated dead cells observed in growth plate chondrocyte layers formed in *Tric-b*-knockout humerus and lib bones. Hematoxylin-eosin–stained tissue sections are presented. Scale bars, 20 *μ*m.

**Figure EV2.**
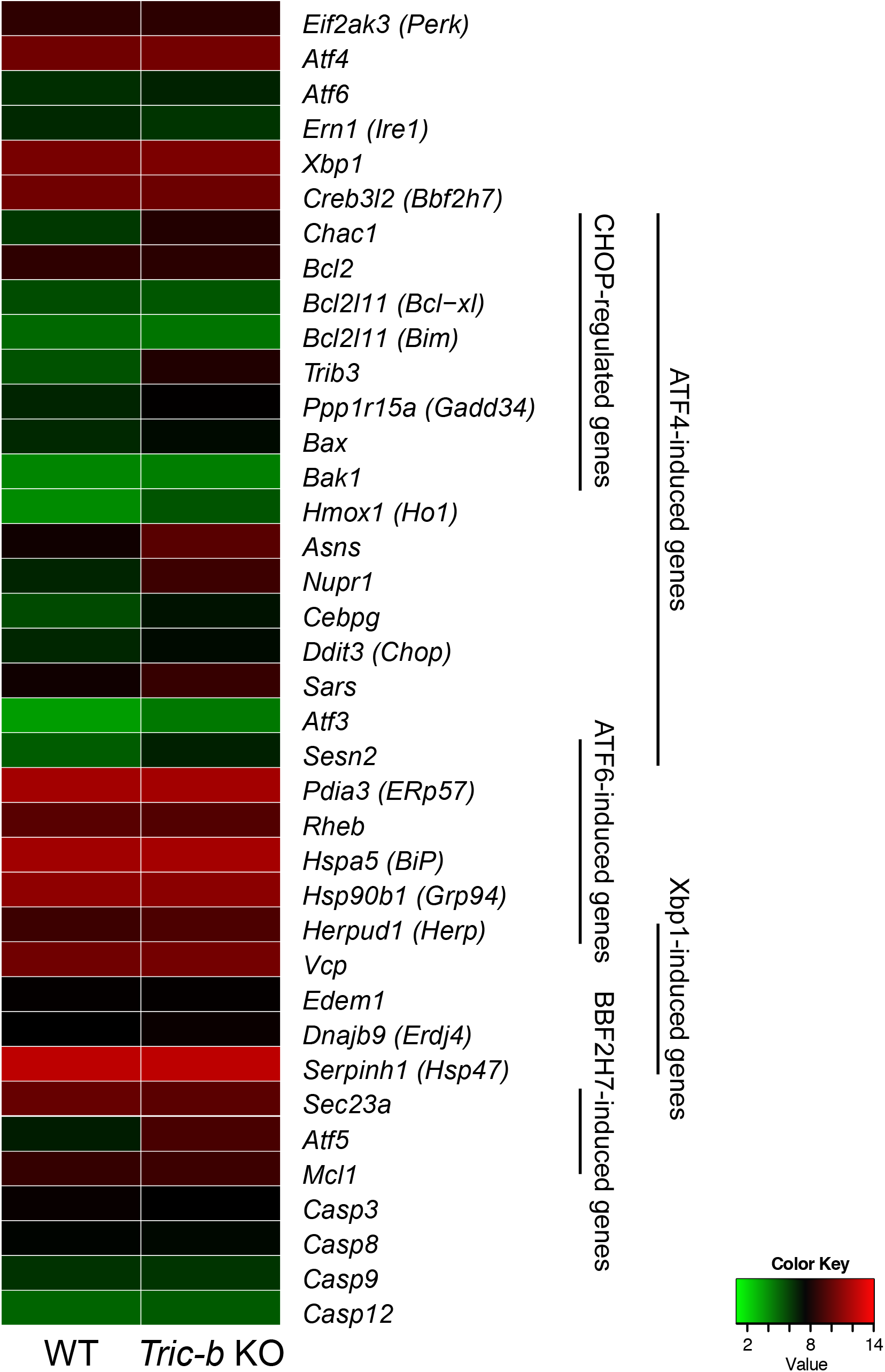
Enhanced expression of PERK-related genes in *Tric-b*-knockout growth plates. Heatmap visually showing UPR-related gene expression in growth plate chondrocytes. Total RNA was prepared from the knockout and wild-type femoral epiphyses enriched in round chondrocytes and subjected to Affymetrix gene-chip analysis. The expression scores yielded are colored according to the color key; green and red denote low and high expression, respectively.

**Table S1.** Antibodies used in this study.

| antigen | immunized animal & conjugation | source & catalogue number |
| --- | --- | --- |
| COL2A1 | Rabbit, Unconjugated | ORIGENE, R1039 |
| KDEL | Mouse, Unconjugated | MBL, M181-3 |
| Mouse IgG (H+L) | Goat, Alexa Fluoro 488 | Thermo Fisher Scientific, A-11029 |
| Rabbit IgG (H+L) | Goat, Alexa Fluo 555 | Thermo Fisher Scientific, A-21429 |
| ATF4 | Mouse, Unconjugated | Santa Cruz Biotechnology, sc-390063 |
| ATF6 | Mouse, Unconjugated | Cosmo Bio, BAM-73-505 |
| BiP | Mouse, Unconjugated | BD Transduction Laboratories, 610979 |
| Cleaved-Caspase-3 | Rabbit, Unconjugated | Cell Signaling Technologies, 9664 |
| Caspase-3 | Rabbit, Unconjugated | Cell Signaling Technologies, 14220 |
| Caspase-8 | Rabbit, Unconjugated | Cell Signaling Technologies, 4927 |
| Cleaved-Caspase-8 | Rabbit, Unconjugated | Novus, NB100-56116 |
| Caspase-9 | Rabbit, Unconjugated | Cell Signaling Technologies, 9504 |
| Caspase-12 | Rabbit, Unconjugated | abcam, ab62484 |
| CHOP | Rabbit, Unconjugated | Cell Signaling Technologies, 5554 |
| Phospho-eIF2α | Rabbit, Unconjugated | Cell Signaling Technologies, 9721 |
| eIF2α | Mouse, Unconjugated | sc-133132 |
| Bbf2h7 | Mouse, Unconjugated | MABE1018 |
| GAPDH | Rabbit, Unconjugated | Sigma-Aldrich, G9545 |
| Rabbit IgG | Mouse, horseradish peroxidase (HRP) | Santa Cruz Biotechnology, sc-2357 |
| mouse IgG | Rabbit, HRP | Dako, P0260 |

**Table S2.**
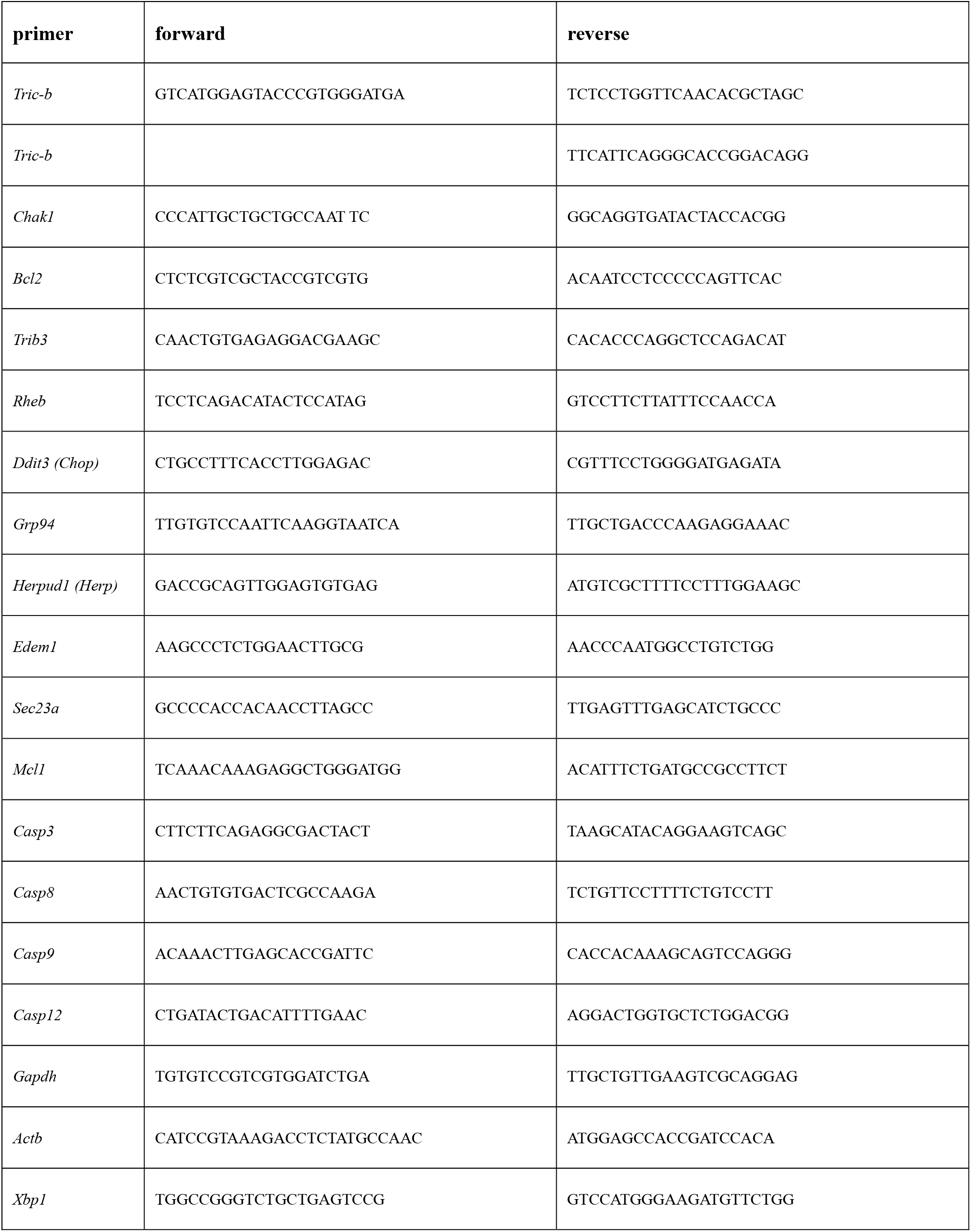
Primers used in this study.

## References

Bateman JF, Cabral WA, Ishikawa M, Garten M, Makareeva EN, Sargent BM, Weis M, Barnes AM, Webb EA, Shaw NJ et al. (2016) Absence of the ER Cation Channel TMEM38B/TRIC-B Disrupts Intracellular Calcium Homeostasis and Dysregulates Collagen Synthesis in Recessive Osteogenesis Imperfecta. PLOS Genetics 12: e1006156

Berendsen AD, Olsen BR (2015) Bone development. Bone 80: 14–18

Byers PH, Pyott SM (2012) Recessively Inherited Forms of Osteogenesis Imperfecta. Annual Review of Genetics 46: 475–497

Coronado R, Miller C (1980) Decamethonium and hexamethonium block K+ channels of sarcoplasmic reticulum Nature 288: 495–497

DuRose JB, Tam AB, Niwa M (2006) Intrinsic capacities of molecular sensors of the unfolded protein response to sense alternate forms of endoplasmic reticulum stress. Mol Biol Cell 17: 3085–3107

Fink RH, Veigel C (1996) Calcium uptake and release modulated by counter-ion conductances in the sarcoplasmic reticulum of skeletal muscle. Acta Physiol Scand 156: 387–396

Gautier L, Cope L, Bolstad BM, Irizarry RA (2004) affy--analysis of Affymetrix GeneChip data at the probe level. Bioinformatics 20: 307–315

Hartl D, Kerbiriou M, Teng L, Benz N, Trouvé P, Férec C (2009) The Calpain, Caspase 12, Caspase 3 Cascade Leading to Apoptosis Is Altered in F508del-CFTR Expressing Cells. PLoS ONE 4: e8436

Hetz C, Zhang K, Kaufman RJ (2020) Mechanisms, regulation and functions of the unfolded protein response. Nature Reviews Molecular Cell Biology 21: 421–438

Ide T, Sakamoto H, Morita T, Taguchi T, Kasai M (1991) Purification of a Cl-channel protein of sarcoplasmic reticulum by assaying the channel activity in the planar lipid bilayer system. Biochem Biophys Res Commun 176: 38–44

Kamp F, Donoso P, Hidalgo C (1998) Changes in luminal pH caused by calcium release in sarcoplasmic reticulum vesicles. Biophys J 74: 290–296

Kasuya G, Hiraizumi M, Maturana AD, Kumazaki K, Fujiwara Y, Liu K, Nakada-Nakura Y, Iwata S, Tsukada K, Komori T et al. (2016) Crystal structures of the TRIC trimeric intracellular cation channel orthologues. Cell Research 26: 1288–1301

Kesavardhana S, Malireddi RKS, Kanneganti T-D (2020) Caspases in Cell Death, Inflammation, and Pyroptosis Annu Rev Immunol 38: 567–295

Kokame K, Kato H, Miyata T (2001) Identification of ERSE-II, a New cis-Acting Element Responsible for the ATF6-dependent Mammalian Unfolded Protein Response. Journal of Biological Chemistry 276: 9199–9205

Lee JK, Kang S, Wang X, Rosales JL, Gao X, Byun H-G, Jin Y, Fu S, Wang J, Lee K-Y (2019) HAP1 loss confers l-asparaginase resistance in ALL by downregulating the calpain-1-Bid-caspase-3/12 pathway. Blood 133: 2222–2232

Liang S-H, Zhang W, McGrath Barbara C, Zhang P, Cavener Douglas R (2005) PERK (eIF2α kinase) is required to activate the stress-activated MAPKs and induce the expression of immediate-early genes upon disruption of ER calcium homoeostasis. Biochemical Journal 393: 201–209

Lupoli TJ, Vaubourgeix J, Burns-Huang K, Gold B (2018) Targeting the Proteostasis Network for Mycobacterial Drug Discovery. ACS Infectious Diseases 4: 478–498

Lv F, Xu X-j, Wang J-y, Liu Y, Asan Wang J-w, Song L-j, Song Y-w, Jiang Y, Wang O et al. (2016) Two novel mutations in TMEM38B result in rare autosomal recessive osteogenesis imperfecta. Journal of Human Genetics 61: 539–545

Meissner G (1983) Monovalent ion and calcium ion fluxes in sarcoplasmic reticulum. Mol Cell Biochem 55: 65–82

Miyazaki Y, Ichimura A, Kitayama R, Okamoto N, Yasue T, Liu F, Kawabe T, Nagatomo H, Ueda Y, Yamauchi I et al. (2022) C-type natriuretic peptide facilitates autonomic Ca2+ entry in growth plate chondrocytes for stimulating bone growth. eLife 11: e71931

Mungrue IN, Pagnon J, Kohannim O, Gargalovic PS, Lusis AJ (2009) CHAC1/MGC4504 is a novel proapoptotic component of the unfolded protein response, downstream of the ATF4-ATF3-CHOP cascade J Immunol 182: 466–176

Pitt SJ, Park K-H, Nishi M, Urashima T, Aoki S, Yamazaki D, Ma J, Takeshima H, Sitsapesan R (2010) Charade of the SR K+-Channel: Two Ion-Channels, TRIC-A and TRIC-B, Masquerade as a Single K+-Channel. Biophysical Journal 99: 417–426

Qian N, Ichimura A, Takei D, Sakaguchi R, Kitani A, Nagaoka R, Tomizawa M, Miyazaki Y, Miyachi H, Numata T et al. (2019) TRPM7 channels mediate spontaneous Ca 2+ fluctuations in growth plate chondrocytes that promote bone development. Sci Signal 12: eaaw4847

Rubinato E, Morgan A, D’Eustacchio A, Pecile V, Gortani G, Gasparini P, Faletra F (2014) A novel deletion mutation involving TMEM38B in a patient with autosomal recessive osteogenesis imperfecta. Gene 545: 290–292

Sadasivan S, Waghray A, Larner SF, Dunn WA, Hayes RL, Wang KKW (2006) Amino acid starvation induced autophagic cell death in PC-12 cells: Evidence for activation of caspase-3 but not calpain-1. Apoptosis 11: 1573–1582

Sadighi Akha AA, Harper JM, Salmon AB, Schroeder BA, Tyra HM, Rutkowski DT, Miller RA (2011) Heightened Induction of Proapoptotic Signals in Response to Endoplasmic Reticulum Stress in Primary Fibroblasts from a Mouse Model of Longevity. Journal of Biological Chemistry 286: 30344–30351

Saito A, Hino S-i, Murakami T, Kanemoto S, Kondo S, Saitoh M, Nishimura R, Yoneda T, Furuichi T, Ikegawa S et al. (2009) Regulation of endoplasmic reticulum stress response by a BBF2H7-mediated Sec23a pathway is essential for chondrogenesis. Nature Cell Biology 11: 1197–1204

Sargeant J, Hay JC (2022) Ca2+ regulation of constitutive vesicle trafficking. Faculty Reviews 11

Shaheen R, Alazami AM, Alshammari MJ, Faqeih E, Alhashmi N, Mousa N, Alsinani A, Ansari S, Alzahrani F, Al-Owain M et al. (2012) Study of autosomal recessive osteogenesis imperfecta in Arabia reveals a novel locus defined byTMEM38Bmutation. Journal of Medical Genetics 49: 630–635

Somlyo AV, Gonzalez-Serratos HG, Shuman H, McClellan G, Somlyo AP (1981) Calcium release and ionic changes in the sarcoplasmic reticulum of tetanized muscle: an electron-probe study. Journal of Cell Biology 90: 577–594

Venturi E, Matyjaszkiewicz A, Pitt SJ, Tsaneva-Atanasova K, Nishi M, Yamazaki D, Takeshima H, Sitsapesan R (2013) TRIC-B channels display labile gating: evidence from the TRIC-A knockout mouse model. Pflugers Arch 465: 1135–1148

Volodarsky M, Markus B, Cohen I, Staretz-Chacham O, Flusser H, Landau D, Shelef I, Langer Y, Birk OS (2013) A Deletion Mutation in TMEM38B Associated with Autosomal Recessive Osteogenesis Imperfecta. Human Mutation 34: 582–586

Wang X-h, Su M, Gao F, Xie W, Zeng Y, Li D-l, Liu X-l, Zhao H, Qin L, Li F et al. (2019) Structural basis for activity of TRIC counter-ion channels in calcium release. Proc Natl Acad Sci 116: 4238–4243

Xu S, Xu Y, Chen L, Fang Q, Song S, Chen J, Teng J (2017) RCN1 suppresses ER stress-induced apoptosis via calcium homeostasis and PERK–CHOP signaling. Oncogenesis 6: e304–e304

Yamazaki D, Komazaki S, Nakanishi H, Mishima A, Nishi M, Yazawa M, Yamazaki T, Taguchi R, Takeshima H (2009) Essential role of the TRIC-B channel in Ca2+ handling of alveolar epithelial cells and in perinatal lung maturation. Development 136: 2355–2361

Yamazaki D, Tabara Y, Kita S, Hanada H, Komazaki S, Naitou D, Mishima A, Nishi M, Yamamura H, Yamamoto S et al. (2011) TRIC-A Channels in Vascular Smooth Muscle Contribute to Blood Pressure Maintenance. Cell Metab 14: 231–241

Yang H, Hu M, Guo J, Ou X, Cai T, Liu Z (2016) Pore architecture of TRIC channels and insights into their gating mechanism. Nature 538: 537–541

Yazawa M, Ferrante C, Feng J, Mio K, Ogura T, Zhang M, Lin P-H, Pan Z, Komazaki S, Kato K et al. (2007) TRIC channels are essential for Ca2+ handling in intracellular stores. Nature 448: 78–82

Zhang Y, Sun R, Geng S, Shan Y, Li X, Fang W (2019) Porcine Circovirus Type 2 Induces ORF3-Independent Mitochondrial Apoptosis via PERK Activation and Elevation of Cytosolic Calcium. Viruses 93: e01784–1718

Zhao C, Ichimura A, Qian N, Iida T, Yamazaki D, Noma N, Asagiri M, Yamamoto K, Komazaki S, Sato C et al. (2016) Mice lacking the intracellular cation channel TRIC-B have compromised collagen production and impaired bone mineralization. Sci Signal 9: ra49

Zhao X, Yamazaki D, Park KH, Komazaki S, Tjondrokoesoemo A, Nishi M, Lin P, Hirata Y, Brotto M, Takeshima H et al. (2010) Ca2+ Overload and Sarcoplasmic Reticulum Instability in tric-a Null Skeletal Muscle. J Biol Chem 285: 37370–37376

Zhou X, Park KH, Yamazaki D, Lin P-h, Nishi M, Ma Z, Qiu L, Murayama T, Zou X, Takeshima H et al. (2020) TRIC-A Channel Maintains Store Calcium Handling by Interacting With Type 2 Ryanodine Receptor in Cardiac Muscle. Circ Res 126: 417–435

Zhu S, McGrath BC, Bai Y, Tang X, Cavener DR (2016) PERK regulates Gq protein-coupled intracellular Ca2+ dynamics in primary cortical neurons. Mol Brain 9: 87

